# Pathogenic Budding Yeasts Isolated outside of Clinical Settings

**DOI:** 10.1101/480251

**Authors:** Dana A. Opulente, Quinn K. Langdon, Kelly V. Buh, Max A. B. Haase, Kayla Sylvester, Ryan V. Moriarty, Martin Jarzyna, Samantha L. Considine, Rachel M. Schneider, Chris Todd Hittinger

**Affiliations:** Laboratory of Genetics, Genome Center of Wisconsin, Wisconsin Energy Institute, J. F. Crow Institute for the Study of Evolution, University of Wisconsin-Madison, Madison, WI 53706; DOE Great Lakes Bioenergy Research Center, University of Wisconsin-Madison, Madison, WI 53706

## Abstract

Budding yeasts are distributed across a wide range of habitats, including as human commensals. However, under some conditions, these commensals can cause superficial, invasive, and even lethal infections. Despite their importance to human health, little is known about the ecology of these opportunistic pathogens, aside from their associations with mammals and clinical environments. During a survey of approximately 1000 non-clinical samples across the United States of America, we isolated 54 strains of budding yeast species considered opportunistic pathogens, including *Candida albicans* and *Candida (Nakaseomyces) glabrata*. We found that, as a group, pathogenic yeasts were positively associated with fruits and soil environments, while the species *Pichia kudriavzevii* (syn. *Candida krusei* syn. *Issatchenkia orientalis*) had a significant association with plants. These results suggest that pathogenic yeast ecology is more complex and diverse than is currently appreciated and raises the possibility that these additional environments could be a point of contact for human infections.

**Importance:** We isolated several opportunistic pathogenic species of yeasts from the subphylum Saccharomycotina from multiple non-clinical environments across the United States of America. Among those strains isolated, over 50% represent the most common opportunistic pathogens. These species, *C. albicans, Candida tropicalis, Candida parapsilosis*, and *C. glabrata*, have been rarely isolated from non-clinical settings and have been usually interpreted as contamination. Our extensive isolations from natural settings challenge this assumption, suggesting that opportunistic pathogens can persist in alternative niches and that their ecology may be more complicated than is currently assumed. Non-clinical environments could be a short-term habitat as these yeasts are passed between their predominant hosts, endothermic animals.

## Observation

Budding yeasts of the subphylum Saccharomycotina are distributed across a wide range of habitats (1–3), including as commensals in the human microbiota (1, 4). However, under rare circumstances, yeasts can cause candidiasis, which is an infection caused by yeasts often assigned to the genus *Candida* and considered opportunistic pathogens (5). While there are rare reports of several species being the agents of candidiasis, including *Saccharomyces cerevisiae* (6), the species *Candida albicans, Candida* (*Nakaseomyces*) *glabrata, Candida parapsilosis*, and *Candida tropicalis* are responsible for approximately 95% of infections (5, 7, 8). Other non-hybrid yeast species recognized as opportunistic pathogens include *Pichia kudriazevii* (syn. *Candida krusei* syn. *Issatchenkia orientalis*), *Candida dublinensis, Candida orthopsilosis, Meyerozyma* (*Candida*) *guilliermondii*, and *Clavispora* (*Candida*) *lusitaniae* (1). *P. kudriavzevii* is the fifth leading cause of yeast infections, is resistant to fluconazole, and has reduced susceptibility to other antifungal treatments (9). Infections by the other species are rare, but each has caused multiple infections in humans (5, 10).

While most are currently classified in the genus *Candida*, these pathogenic species belong to several phylogenetically distinct clades (5). Despite this diversity, one commonality is that little is known about their ecology; it is unclear whether their primary ecological niche is endothermic animals and clinical settings, or whether they are adapted to additional environments. While disease-causing species have been recovered from a variety of habitats, such as food and clothing, these isolates are generally interpreted as contamination, rather than being sourced from a yeast habitat (1, 11–13).

Understanding the ecology of pathogenic yeasts is critical to human health for multiple reasons. First, mortality from infections by these yeasts remains high, and candidiasis is the fourth most common hospital-associated bloodstream infection (4, 14, 15). Second, *Candida auris* recently emerged as a multi-drug resistant opportunistic pathogen, with additional strains exhibiting resistance to many anti-fungal drugs (14, 16). Finally, it is unclear whether there might be environmental reservoirs that could act as contact points for their primary hosts, or whether the niches of pathogenic yeasts are indeed exclusively endothermic animals and clinical environments (17).

We extensively sampled non-clinical substrates across the United States of America to enrich for and isolate yeasts. In total, we collected approximately 1000 samples and have isolated about 5000 strains of yeasts. Across our sampling regime, we isolated 54 strains of species that are considered opportunistic pathogens (Figure 1, Table S1). Recently and independently, three strains of *C. albicans* were isolated from oak bark in New Forest in the United Kingdom (13). Collectively, these results suggest that pathogenic yeast ecology is more complex and diverse than is currently appreciated and raises the possibility that these additional environments could be a point of contact for infections.

**Figure 1:**
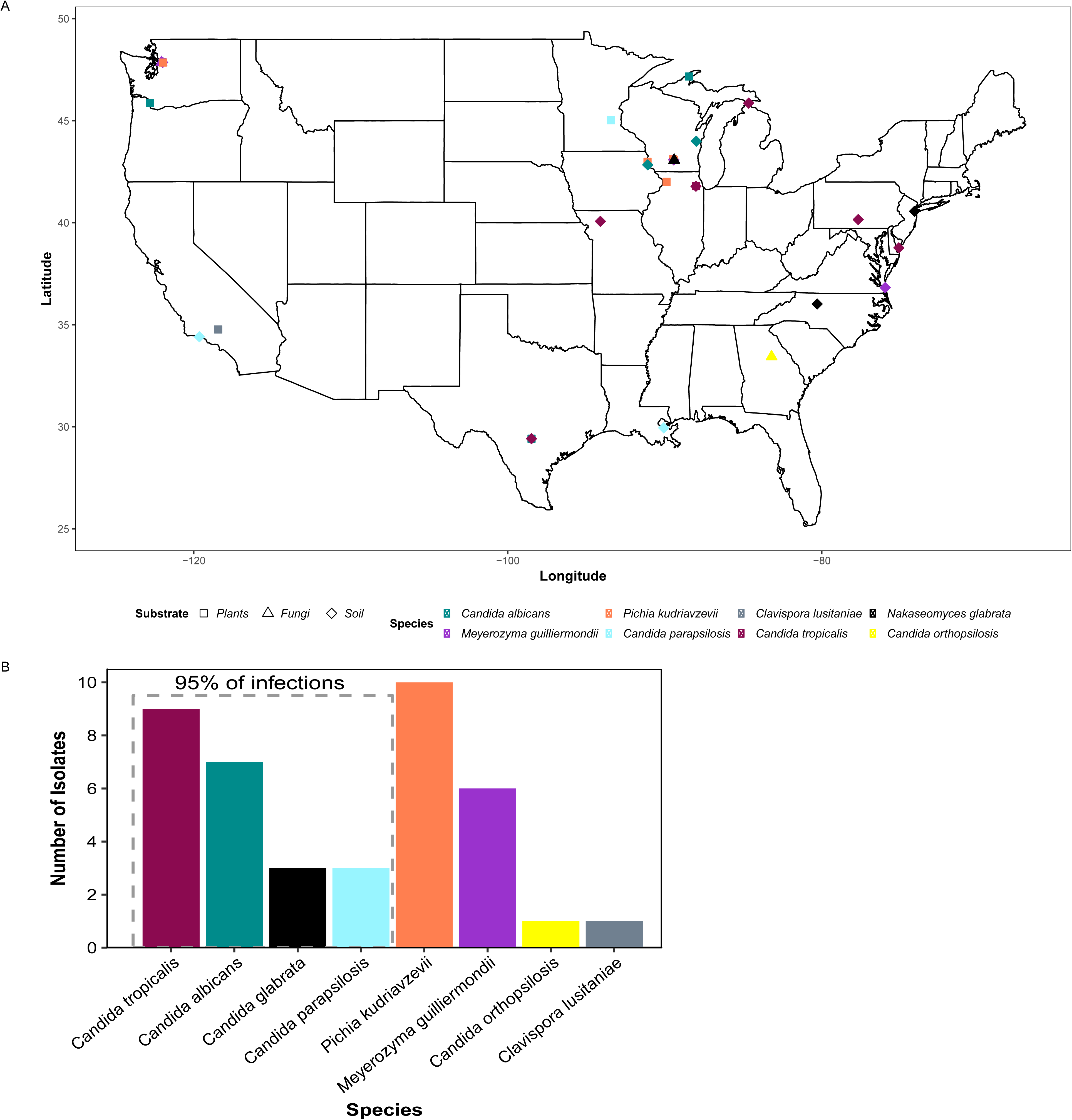
Opportunistic pathogenic species of yeasts in the subphylum Saccharomycotina are found throughout the United States of America on a variety of non-clinical substrates. A) Map displaying where opportunistic pathogens were isolated. The colors represent the species isolated, and the shapes of the points represent the substrate from which they were isolated. B) Bar graph of unique strains of each species.

The strains we isolated came from 40 different samples; in 14 cases, multiple strains of a single species were isolated from a single sample, often from different isolation temperatures (Table S1). Once those were collapsed, we had 40 unique strains, 55% of which were from the four species that cause 95% of candidiasis infections (Figure 2). The species isolated most was *P. kudriavzevii*, making up 25% (n = 10) of our isolates of opportunistic pathogenic species that were isolated from non-clinical settings several times across a wide range of environments, including fermentations, soil, and fruits (1, 18). Its prevalence in our environmental isolations is consistent with previous studies.

**Figure 2:**
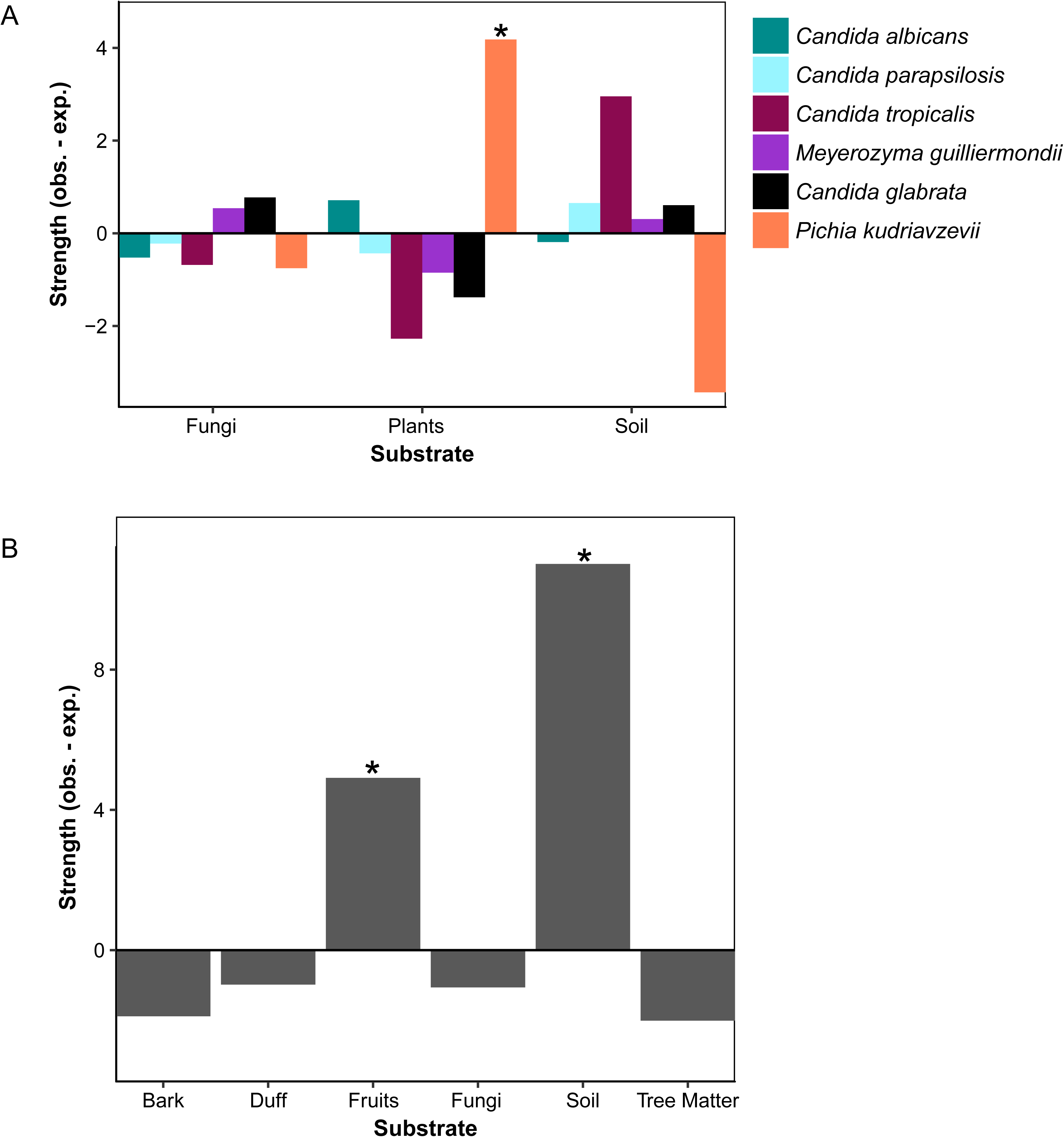
Pathogenic yeast species are associated with non-clinical/non-endothermic environments. A) *P. kudriavzevii* was a significantly associated with plants (p_adj._ = 0.018). Bar graphs representing the strength (difference between observed and expected counts) of associations between pathogenic species and isolation substrates. B) Opportunistic pathogens were associated with soil (p_adj._ < 0.001) and fruits (p_adj._ = 0.021). Bar graph representing the strength of association between pathogenic yeasts and specific isolation environments.

While *P. kudriazevii* and *M. guilliermondii* have been isolated from non-clinical/ectothermic environments (19, 20), the other species we isolated have only been rarely isolated from non-clinical settings. It is unknown whether these species are actively growing in these environments, but these non-clinical environments could potentially act as a secondary niche for these species. Furthermore, these environments could be an additional source of contact between these species and their primary hosts, potentially as the organism passes between hosts. To determine whether pathogenic yeasts were associated with particular non-clinical environments, we classified our samples in two ways. We used a more general description of whether it was isolated from plants, soil, or fungi; and then, when the specific information was available, a more specific description of the substrate (e.g. fruits, leaves, sand) was given. This method was used to determine whether specific parts of a substrate were more important for pathogenic yeasts than others.

Samples varied significantly in terms of the number of species isolated from an environment (Table S1). Across all species with multiple isolation events, species were not limited to a single substrate. For example, *C. albicans* was isolated from fruits, soil, and plant matter. *P. kudriazevii* was significantly associated with isolation from plants (p_adj_ = 0.0185) (Figure 2A, Table S4). We detected no other significant associations among substrates at the general descriptive level for the other species in our data.

Due to the smaller sample sizes at the specific descriptive levels for our substrates, we did not test for associations between them and species. However, at this level, our opportunistic pathogen isolates collectively exhibited significant associations with fruits (p_adj._ = 0.021) and soil (p_adj._ < 0.001) (Figure 2B, Table S5). The association between fruits and our isolates was most likely driven by our *P. kudriavzevii* isolates, which were, in our more general analysis, positively associated with plants. This association could be the result of animals acting as a vector to transport these opportunistic pathogenic species to fruits they regularly visit. The association with soil was mostly driven by *C. tropicalis*, but other species also contributed; 39% (n = 7) of our soil isolates were *C. tropicalis*. Our results, combined with the independent isolation of three strains of *C. albicans* from oaks in the United Kingdom (13), suggest that these alternative niches could potentially act as a secondary contact site between these pathogenic yeasts and their human hosts.

### Methods of sample collection, yeast isolation, quality control, and statistical analyses

Yeasts were collected throughout the United States from multiple substrates by members of the Hittinger Lab, as well as citizen scientists (Table S1). Statistical analyses did not detect bias in pathogenic isolates coming from any individual collector (χ^2^ = 0.072, p-value = 1, Table S2) or among individuals isolating yeasts from samples (χ^2^ = 1.63, p-value = 1, Table S3). All samples were collected using sterile bags and did not come into contact with the humans collecting them. Samples were processed by different members of the lab using published yeast enrichment and isolation protocols (21). Negative controls were included to ensure no contamination occurred. After enrichment, yeast species were identified by ITS sequencing, as previously described (21).

We removed strains from our analyses that were isolated from areas that are highly trafficked by humans, such as samples that were isolated from compost piles. We collapsed multiple strains isolated from the same sample (including when isolated at different temperatures) to one representative strain since the strains may be closely related. We determined positive associations using modified methods previously described (3). All analyses were done in the statistical language R (v. 3.3.0). All graphs were made using the R package *ggplot2* (v. 2.2.1).

## Acknowledgments

We thank the following citizen scientists or lab members for collecting samples or generating preliminary results: Aaron G. Barton, Amanda Beth Hulfachor, Russell L. Wrobel, Leslie Shown, The Zasadil Family, EmilyClare P. Baker, Drew T. Doering, Angela Sheddan, Sarah Wright, Bill Saucier, Bill Vagt, Annette Opulente, and Sophie Smead. This material is based upon work supported by the National Science Foundation under Grant Nos. DEB-1253634 (to CTH), DEB-1442148 (to CTH), and DGE-1256259 (Graduate Research Fellowship to QKL); funded in part by Lakeshore Nature Preserve Student Engagement Grants (to MJ and RMS); and funded in part by the DOE Great Lakes Bioenergy Research Center (DOE Office of Science BER DE-SC0018409 to Timothy J. Donohue). QKL was also supported by the Predoctoral Training Program in Genetics, funded by the National Institutes of Health (5T32GM007133). CTH is a Pew Scholar in the Biomedical Sciences and Vilas Faculty Early Career Investigator, supported by the Pew Charitable Trusts and Vilas Trust Estate, respectively.

## Supplementary Table

**Table S1:** All opportunistic pathogens isolated across the United States of America, including the GPS location, substrate, isolation temperature, sample ID, and the ITS and/or D1/D2 sequence used to confirm species identification.

**Table S2:** Number of samples collected by individuals whose samples contained an opportunistic pathogenic species. The “RG2 prop” column represents the proportion of opportunistic pathogens obtained from samples collected by that individuals normalized by the total samples collected by the individual.

**Table S3:** Number of samples isolated by individuals who isolated an opportunistic pathogenic species. The “RG2 prop” column represents the proportion of opportunistic pathogens isolated by that individuals normalized by the total number of isolates obtained by the individual.

**Table S4:** Associations among species and general isolation substrates. The expected value is the average value from the randomized data (n = 1000).

**Table S5:** Associations among pathogenic yeasts and specific isolation substrates. The expected value is the average value from the randomized data (n = 1000).

## References

1. Kurtzman C, Fell JW, Boekhout T. 2011. The yeasts: A taxonomic study. Elsevier.

2. Buzzini P, Lachance M-A, Yurkov A. 2017. Yeasts in natural ecosystems: diversity. Springer.

3. Opulente DA, Rollinson EJ, Bernick-Roehr C, Hulfachor AB, Rokas A, Kurtzman CP, Hittinger CT. 2018. Factors driving metabolic diversity in the budding yeast subphylum. BMC Biol 16:26.

4. Gabaldón T, Fairhead C. 2018. Genomes shed light on the secret life of *Candida glabrata*: not so asexual, not so commensal. Curr Genet 1–6.

5. Gabaldón T, Naranjo-Ortíz MA, Marcet-Houben M. 2016. Evolutionary genomics of yeast pathogens in the Saccharomycotina. FEMS Yeast Res 16:fow064.

6. Strope PK, Skelly DA, Kozmin SG, Mahadevan G, Stone EA, Magwene PM, Dietrich FS, McCusker JH. 2015. The 100-genomes strains, an *S. cerevisiae* resource that illuminates its natural phenotypic and genotypic variation and emergence as an opportunistic pathogen. Genome Res 25:762–774.

7. Pfaller MA, Diekema DJ. 2007. Epidemiology of Invasive Candidiasis: a Persistent Public Health Problem. Clin Microbiol Rev 20:133–163.

8. Diekema D, Arbefeville S, Boyken L, Kroeger J, Pfaller M. 2012. The changing epidemiology of healthcare-associated candidemia over three decades. Diagn Microbiol Infect Dis 73:45–48.

9. Pelletier R, Alarie I, Lagacé R, Walsh TJ. 2005. Emergence of disseminated candidiasis caused by *Candida krusei* during treatment with caspofungin: case report and review of literature. Med Mycol 43:559–564.

10. Pfaller MA, Diekema DJ, Gibbs DL, Newell VA, Ellis D, Tullio V, Rodloff A, Fu W, Ling TA, Group GAS. 2010. Results from the ARTEMIS DISK Global Antifungal Surveillance Study, 1997 to 2007: a 10.5-year analysis of susceptibilities of *Candida* species to fluconazole and voriconazole as determined by CLSI standardized disk diffusion. J Clin Microbiol 48:1366–1377.

11. Van Uden N, Faia MDEM, Assis-Lopes L. 1956. Isolation of *Candida albicans* from vegetable sources. Microbiology 15:151–153.

12. Di Menna ME. 1958. *Candida albicans* from grass leaves. Nature 181:1287.

13. Bensasson D, Dicks J, Ludwig JM, Bond CJ, Elliston A, Roberts IN, James SA. 2018. Diverse lineages of Candida albicans live on old oaks. bioRxiv. doi: https://doi.org/10.1101/341032

14. Revie NM, Iyer KR, Robbins N, Cowen LE. 2018. Antifungal drug resistance: evolution, mechanisms and impact. Curr Opin Microbiol 45:70–76.

15. Morgan J, Meltzer MI, Plikaytis BD, Sofair AN, Huie-White S, Wilcox S, Harrison LH, Seaberg EC, Hajjeh RA, Teutsch SM. 2005. Excess mortality, hospital stay, and cost due to candidemia: a case-control study using data from population-based candidemia surveillance. Infect Control Hosp Epidemiol 26:540–547.

16. Dolande M, García N, Capote AM, Panizo MM, Ferrara G, Alarcón V. 2017. *Candida auris*: Antifungal Multi-Resistant Emerging Yeast. Curr Fungal Infect Rep 11:197–202.

17. Daszak P, Cunningham AA, Hyatt AD. 2000. Emerging Infectious Diseases of Wildlife--Threats to Biodiversity and Human Health. Science 287:443 LP–449.

18. Douglass AP, Offei B, Braun-Galleani S, Coughlan AY, Martos AAR, Ortiz-Merino RA, Byrne KP, Wolfe KH. 2018. Population genomics shows no distinction between pathogenic *Candida krusei* and environmental *Pichia kudriavzevii*: One species, four names. PLOS Pathog 14:e1007138.

19. Brandão LR, Vaz ABM, Santo LCE, Pimenta RS, Morais PB, Libkind D, Rosa LH, Rosa CA. 2017. Diversity and biogeographical patterns of yeast communities in Antarctic, Patagonian and tropical lakes. Fungal Ecol 28:33–43.

20. Moubasher AH, Abdel-Sater MA, Zeinab SM. 2018. Diversity of floricolous yeasts and filamentous fungi of some ornamental and edible fruit plants in Assiut area, Egypt. Curr Res Environ Appl Mycol 8:135–161.

21. Sylvester K, Wang Q-M, James B, Mendez R, Hulfachor AB, Hittinger CT. 2015. Temperature and host preferences drive the diversification of *Saccharomyces* and other yeasts: A survey and the discovery of eight new yeast species. FEMS Yeast Res 15:fov002.

